# A comprehensive personal omics clinical interpreter based on genomic and transcriptomic profiles

**DOI:** 10.1101/2024.02.22.581482

**Authors:** Yaqing Liu, Qingwang Chen, Qiaochu Chen, Leqing Sang, Yunjin Wang, Leming Shi, Yuanting Zheng, Li Guo, Ying Yu

**Affiliations:** State Key Laboratory of Genetic Engineering, Human Phenome Institute, School of Life Sciences, and Shanghai Cancer Center, Fudan University, Shanghai, China; The International Human Phenome Institutes, Shanghai, China; Department of Breast Surgery, Precision Cancer Medicine Center, Key Laboratory of Breast Cancer in Shanghai, Fudan University Shanghai Cancer Center, Shanghai, China; Department of Oncology, Shanghai Medical College, Fudan University, Shanghai, China; State Key Laboratory of Multiphase Complex Systems, Institute of Process Engineering, Chinese Academy of Sciences, Beijing, China; School of Chemical Engineering, University of Chinese Academy of Sciences, Beijing, China

**Keywords:** Precision medicine, Pharmacogenomics, Therapeutic interpretations, Drug prioritization, Drug response, Integrative genomic profiling

## Abstract

Advances in precision medicine rely on the accurate identification and analysis of molecular alterations for personalized diagnostic, prognostic, and therapeutic decision-making. A critical obstacle is the integration of heterogeneous interpretations of clinically actionable alterations from various knowledgebases. Here, we present the Personal Omics Interpreter (POI), a web-based application engineered to aggregate and interpret therapeutic options, including targeted, immunological, and chemotherapeutic agents, by leveraging personal genomic and transcriptomic profiles. POI employs the Precision Medicine Knowledgebase (PreMedKB), an updated harmonized resource we previously reported, to annotate the clinically actionable somatic variants. It further incorporates a predictive algorithm to broaden therapeutic options according to established gene-gene interactions and offers insights into phenotypic responses of chemotherapeutic agents through phasing germline diplotypes. Validated against three cohort datasets encompassing over 22,000 cancer patients, POI demonstrates consistently high matching rates (94.7 ∼ 95.6%) between patients and suggested therapies, highlighting its potential in supporting precision-driven informed treatment strategies.

## Background

Precision medicine represents a paradigm shift in healthcare, offering a new approach to optimize treatment outcomes through the customization of therapeutic interventions according to the patient’s unique molecular profiles [1]. This shift aims to maximize efficacy and minimize the occurrence of adverse drug reactions [2]. As the application of omics data becomes increasingly prevalent in this domain, there arises a growing demand for identifying and interpreting clinically actionable alterations across both scientific research and clinical domains [3–5]. This pressing demand stems from the desire to effectively prioritize anti-cancer drugs based on a comprehensive understanding of the molecular landscape, ensuring targeted and precise treatment strategies for patients.

Several interpretation tools have been developed to address this demand, primarily focusing on somatic alterations [6–11]. However, a select few, such as PORI [12], MOAlmanac [13], CCAS [14], and PanDrugs2.0 [7, 15], have broadened their interpretative scope to encompass germline variants, RNA outliers, and other relevant factors. These platforms employ diverse strategies to analyze genomic and transcriptomic characteristics, underscoring the significant potential of multi-dimensional data interpretation in identifying actionable therapeutic alterations. Despite these advancements, existing platforms still exhibit certain limitations, including incomplete coverage of interpreted data types (e.g., RNA expression and genotype data), limited exploration of cross-omics features, and constrained capabilities in therapeutic recommendations, such as chemotherapy, beyond targeted therapies [12, 16, 17]. Thus, achieving an accurate interpretation of multi-dimensional molecular changes remains a substantial challenge in advancing precision medicine.

The gap for a comprehensive platform that integrates genomic and transcriptomic data to adeptly prioritize anti-cancer drugs individually is evident. First of all, exploring gene expression-based inference of cancer drug sensitivity has emerged as a promising avenue for identifying actionable therapeutic alterations based on RNA expression data [18]. Additionally, accumulating evidence suggests that targeting co-occurring oncogenic driver aberrations holds promise for robust and durable therapeutic responses, emphasizing the significance of pathway analysis in interpreting actionable therapeutic alterations [19]. Furthermore, the impact of germline variants and their genotypes on the efficacy, dosage, and toxicity of conventional chemotherapy has been recognized, further highlighting their relevance in identifying actionable therapeutic alterations [20].

Addressing this imperative, we develop the Personal Omics Interpreter (POI), a user-centric tool that utilizes a multiomics integrative strategy to identify clinically actionable alterations for anti-cancer drug prioritization (https://premedkb.cn/poi/#/). POI is designed to accommodate multi-dimensional alterations as input, including somatic and germline single nucleotide variations (SNVs), small insertions and deletions (Indels), copy number variations (CNVs), gene fusions, tumor mutational burden (TMB) and microsatellite instability (MSI), pathogenic germline variants, and aberrantly expressed genes. POI also employs a predictive algorithm that enables the inference of suitable drugs for patients lacking straightforward actionable therapeutic alterations. Extensive validation testing of POI has been conducted using prominent datasets, including the Cancer Genome Atlas (TCGA) multi-cancer datasets and the MSK-IMPACT datasets, as well as our proprietary breast cancer dataset. These results underscore the effectiveness and reliability of POI in aiding precision medicine decision-making and prioritizing anti-cancer drugs across various cancer types.

## Results

### Architecture of POI

POI is a comprehensive clinical interpretation algorithm designed to facilitate the integrated interpretation of genomics and transcriptomics data to prioritize drugs for individual cancer patients. The architecture of POI is depicted in **Fig. 1**. POI consists of four key components: (1) a backend knowledgebase, named PreMedKB, which serves as a comprehensive data repository comprising information on the “gene-variant-disease-drug” model to facilitate comprehensive interpretation; (2) multiomics profile as input: POI effectively deciphers the genomics variants (somatic and germline SNVs/Indels, CNV, and fusion), genomics signatures (TMB, MSI), and aberrantly expressed genes (transcriptomics alterations) of a patient to prioritize targeted and immunological drugs; (3) modules designed to perform multiple tasks, including the parsing of multiomics profiles, identification of actionable alterations, inference of off-label drugs, and interpretation using harmonized evidence; (4) a user-friendly web interface that generates therapeutic reports of therapeutic interpretations.

**Fig. 1.**
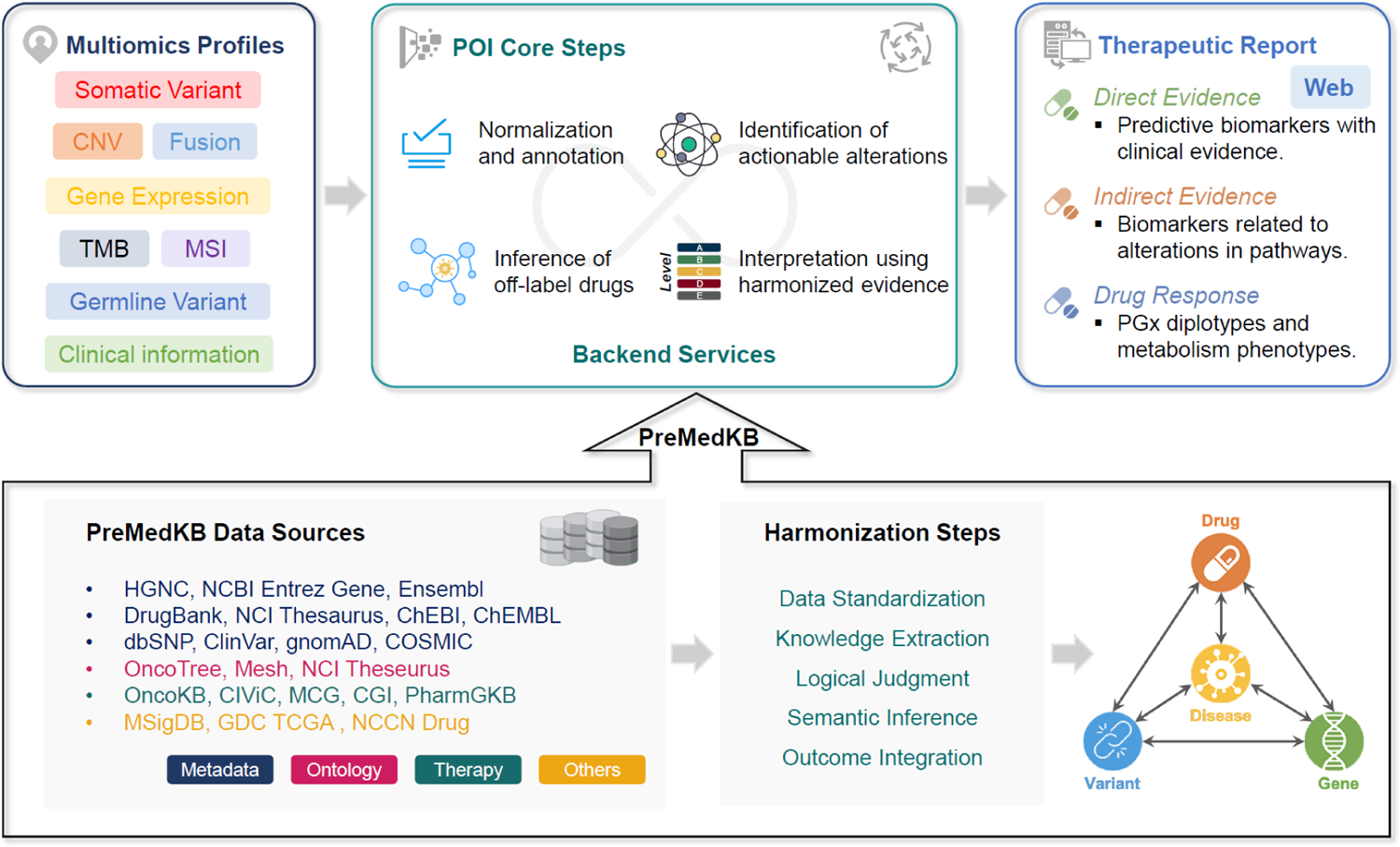
Architecture of POI. POI consists of three key components: (1) a backend knowledgebase, PreMedKB, which serves as a comprehensive data repository comprising information on the “gene-variant-disease-drug” model to facilitate comprehensive interpretation; (2) specialized modules (POI core steps) designed to perform essential tasks, including the parsing of multiomics profiles, identification of actionable alterations, and interpretation based on harmonized evidence; and (3) a user-friendly web interface that generates therapeutic reports for prioritizing anti-cancer drugs.

The overall process of how POI works is shown in **Fig. 2**. Briefly, POI utilizes a harmonized knowledgebase to perform a comprehensive analysis of multiple feature sets within the patient’s multiomics profiles. POI can provide three classes of clinical evidences for comprehensive annotation of drug biomarkers based on variation types (somatic or germline variants) and clinical confidences (direct evidence with high confidence or predictive evidence with lower confidence). The clinical evidences include (1) direct evidence: druggable biomarkers with direct clinical evidence; (2) indirect evidence: predictive biomarkers that are found to be involved in cancer-specific pathway(s) and interacted with druggable biomarkers; and (3) drug response: prediction of patient sensitivity to chemotherapeutic drugs based on germline variations and/or combination of germline variations within specific genes (gene haplotypes or diplotypes). After analyzing in POI, the generated report can be obtained, covering essential information such as drug prioritization, actionable alterations, and drug response prediction based on pharmacogenomic replicates and metabolic phenotypes.

**Fig. 2.**
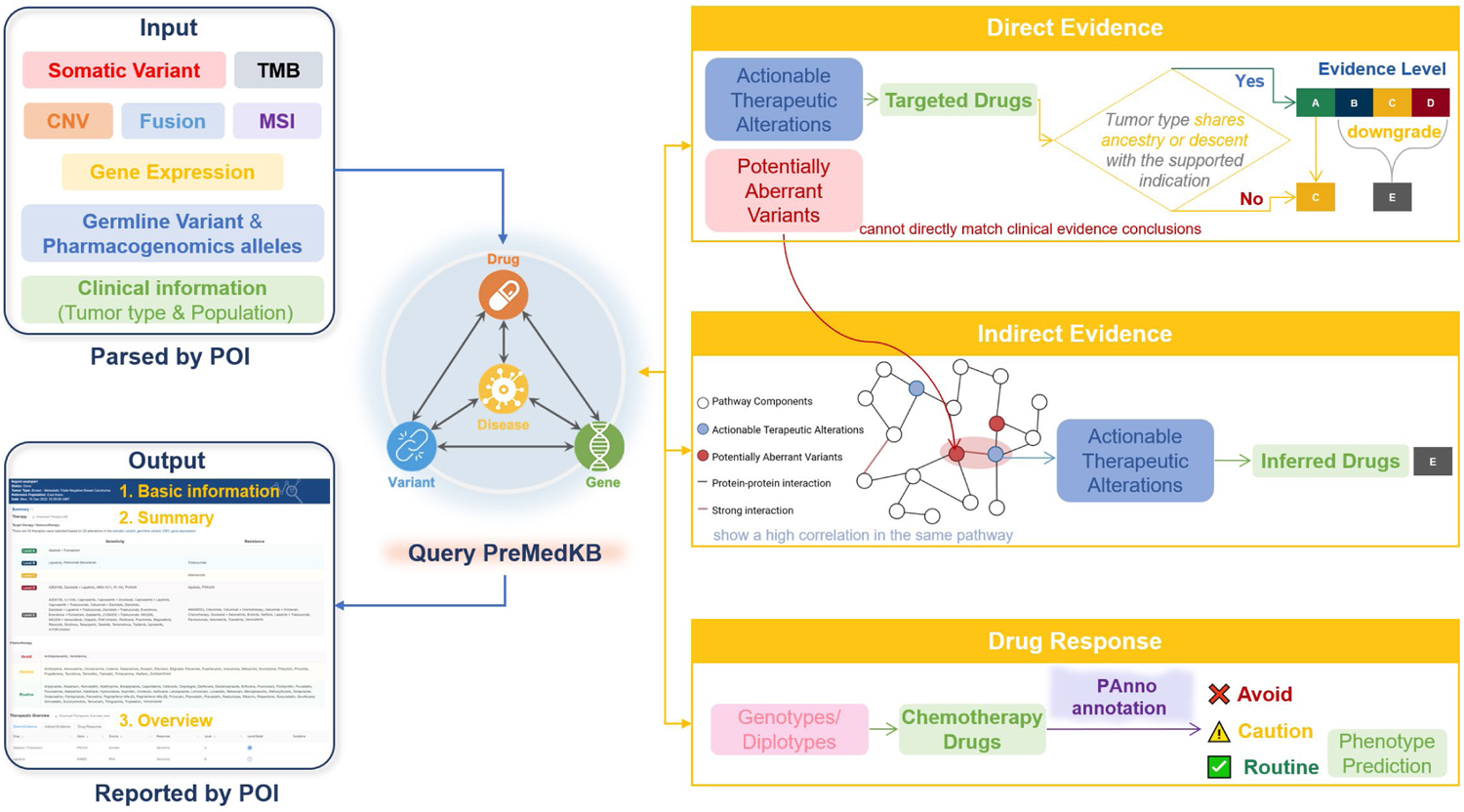
Flowchart of POI core steps. POI employs a comprehensive analysis of somatic and germline SNV/Indel, CNV, gene fusions, TMB, MSI, and patient gene expression files to identify targeted and chemotherapy drugs. Direct evidence involves precise matching of variants with PreMedKB entries, leading to the identification of actionable therapeutic alterations. The assigned grade (A, B, C, or D) for drug prioritization depends on whether the patient’s tumor type shares ancestry or descent with the supported indication in the clinical evidence. If not, the assigned grade is downgraded accordingly (level A to level C, other levels to level E). Indirect evidence relies on the identification of potential aberrant variants and the assessment of associated actionable therapeutic alterations within the same biological pathway. Inferred drugs in this context are assigned grade E. Furthermore, drug response analysis involves resolving germline genotypes/diplotypes and predicting patient phenotypes based on relevant pharmacogenomic alleles. The drugs are classified into three categories based on their recommended use: avoid, caution, and routine.

### Comprehensive data integration and normalization of PreMedKB

PreMedKB encompassed a comprehensive collection of cancer therapy data, including 502 diseases, 458 genes, 6,713 variants, and 865 drugs. The semantic network within PreMedKB revealed numerous associations, such as 6,713 gene-variant associations, 2,493 gene-disease associations, 3,168 gene-drug associations, 41,168 variant-drug associations, 51,316 variant-disease associations, and 4,777 drug-disease associations (**Fig. 3a**). Notably, the involvement of various tissues in the support analysis conducted by POI leads to substantial variation in the number of diseases, drug-disease associations, gene-disease associations, and variant-disease associations across different tissues (**Fig. S1**). Additionally, PreMedKB focused on diverse variants, including 4,840 SNVs, 733 Indels, 247 fusions, 212 gene expressions, 128 CNVs, 57 haplotypes, 38 structural variations (SVs), 5 genomic signatures, and 522 other variant types (**Fig. 3b**). The rich data coverage expanded the breadth and depth of the knowledgebase, enabling it to provide comprehensive, and accurate information, thereby enhancing the comprehensiveness of the knowledgebase.

**Fig. 3.**
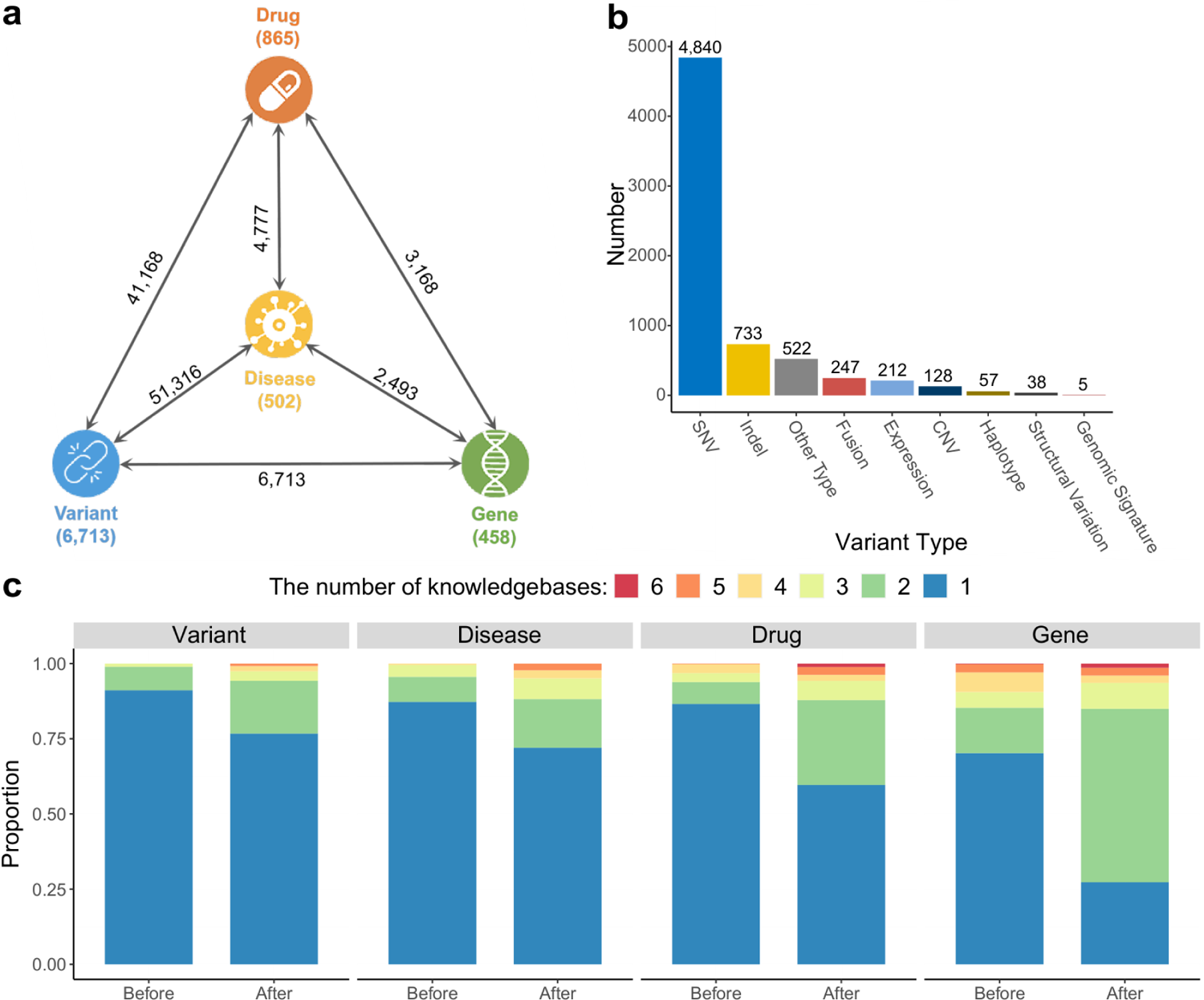
Comprehensive data integration and normalization of PreMedKB. **a** Relationships between multiple diseases, genes, variants, and drugs, emphasizing their relevance to tumor therapy. **b** Bar plots display the number of actionable therapeutic alterations categorized accordingly. **c** The comparison of element uniqueness across knowledgebases before and after normalization, respectively.

To facilitate the assessment of aberrant gene expression, a comprehensive RNA reference database was constructed using gene expression data from diverse cancer types within the TCGA RNA expression landscape. The utilization of this reference database on a web server allowed for the evaluation of patients’ gene expression levels, providing insights into the gene expression distribution specific to their cancer type (**Fig. S2a**). Additionally, **Fig. S2b** and **Fig. S2c** illustrated the expression distribution of gene *TP53* across different cancer types and the expression distribution of five key genes associated with breast cancer, respectively. These results exemplified the comprehensive gene expression data coverage in the reference database, allowing for a deeper understanding of gene expression patterns across different cancers and within specific cancer types.

When integrating the knowledge from different authoritative precision medicine databases, heterogeneity exists due to their distinctive structures and contents [11]. To address this, knowledge normalization techniques were employed to eliminate redundancy, enhance data interoperability, establish consistent data standards, and improve data integration capabilities. These efforts aimed to provide users with more comprehensive, accurate, and reliable information from the knowledgebase. Analyzing the four elements individually revealed a significant overlap in database construction (**Fig. 3c**), however, approximately 91% of variants, 88% of diseases, 87% of drugs, and 70% of genes remained unique across knowledgebases. Through the normalization of metadata terminologies, we identified mutually interpretable terms among the knowledgebases, leading to a reduction in uniqueness to 77% for variants, 72% for diseases, 60% for drugs, and 27% for genes, which decreased the heterogeneity of knowledge in the PreMedKB.

By expanding data coverage and implementing knowledge normalization, the precision, and comprehensiveness of the knowledgebase is greatly improved. This contributes to the comprehensive identification and interpretation of actionable therapeutic alterations, providing valuable insights for precision medicine applications.

### Validation of the effectiveness of POI based on cohort datasets

To assess the effectiveness of POI for comprehensive precision drug prioritization based on multiomics data of individual patients, we conducted a thorough evaluation using three cohort datasets: TCGA [21], MSK-IMPACT [22], and Triple-negative Breast Cancer of Fudan Shanghai Cancer Center (FUSCC) cohort [23, 24] (**Fig. S3**).

The TCGA dataset included somatic genomic and transcriptomic information for six categories of patients: somatic variants (10,030 patients), CNV (10,667 patients), fusion (6,306 patients), TMB (1,043 patients), MSI (422 patients), and gene expression (702 patients), covering a wide range of tumor types, including primary and metastatic cases. Widely recognized for its extensive sample size and diverse data types, the TCGA dataset served as a cornerstone in cancer research and evaluation. The MSK-IMPACT dataset, generated by Memorial Sloan Kettering Cancer Center (MSKCC), consisted of somatic genomic information for five groups of patients: somatic variants (including SNV and Indels) (10,129 patients), CNV (10,945 patients), fusion gene (1,171 patients), MSI (180 patients), and TMB (988 patients). The MSK-IMPACT dataset was pivotal in clinical oncology, representing a widely utilized resource for understanding metastatic cancer genomics. Finally, the FUSCC dataset, centered on a single tumor type (triple-negative breast cancer, TNBC), offered a rich resource of integrative genomic information, including both somatic and germline genomic data, as well as transcriptomic profiles. The FUSCC dataset can be split into six categories of test files: somatic variants (279 patients), germline variants (279 patients), CNV (401 patients), TMB (57 patients), and gene expression (88 patients). Haplotype/diplotype information and corresponding drug responses were obtained based on germline variants using the *PAnno* tool [25] in the drug response module. Therefore, the inclusion of these datasets ensures comprehensive validation of the performance of POI.

The validation results demonstrated that POI was able to match at least one drug for approximately 95% of patients across the three cohorts (**Fig. 4a**), where over 39.4% of patients can obtain the reliable drug prioritizations of Level A and Level B. In the TCGA cohort, POI exhibited comparable performance to the latest platform, PORI [12], with drug prioritizations available for approximately 96% of patients. Additionally, in the MSK-IMPACT cohort, POI significantly improved the prioritized drug ratios compared to the original report from MSKCC [26], providing drug prioritizations for approximately 94.7% of patients. The distribution of the highest drug evidence levels in each tissue within the MSK-IMPACT cohort mirrored the results observed in the TCGA cohort. These findings highlight the expanding knowledgebases of POI and its ability to prioritize drugs using pathway inference strategies, even in off-label use cases where no previous drugs were available.

**Fig. 4.**
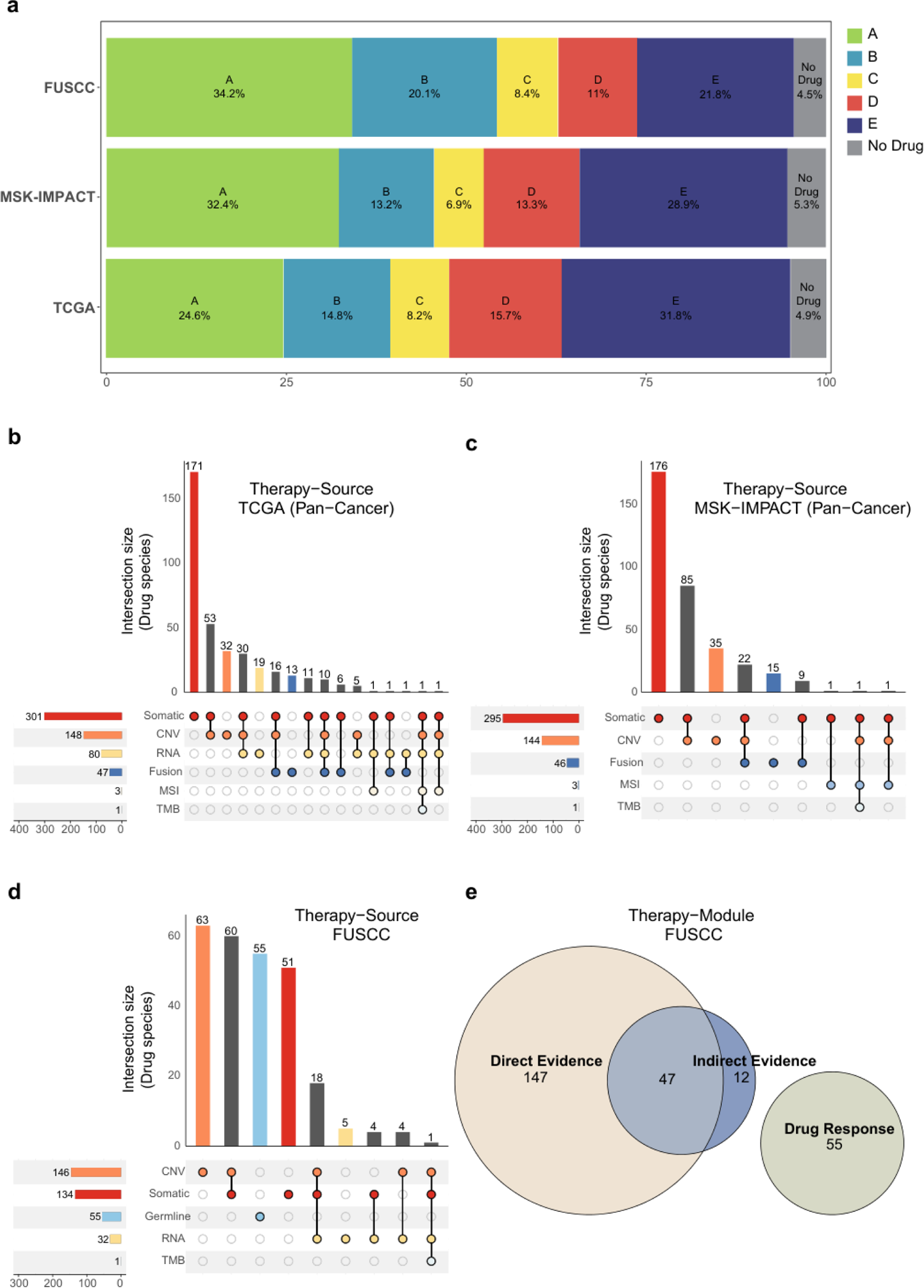
Validation of precision drug prioritization on three external cohorts. Performance validation of precision drug prioritization based on multiomics data from three external cohorts, namely TCGA, MSK-IMPACT, and FUSCC cohort. **a** Barplot of the distribution of highest levels of drugs that patients received from POI in the three cohorts, including level A, B, C, D and E with no drug. **b**-**d** Upset plots of the number of recommended drugs for TCGA MSK-IMPACT, and FUSCC cohorts, respectively. Variants are divided into seven types: single nucleotide variants and indels from somatic mutation (Somatic), copy number variants (CNV), gene fusion (Fusion), high microsatellite instability (MSI), high tumor mutation burden (TMB), RNA gene expression (RNA) variants, and genotype from germline mutation (Germline). Side bar plots represent the aggregate drug species matched to specific variant categories, while top bar plots indicate the count of drug species within each intersection group. **e** The Venn diagram displays the number of recommended drug species from different modules of POI in FUSCC cohort.

Though somatic and CNV input files predominantly contributed to drug prioritizations in the TCGA, MSK-IMPACT, and FUSCC cohorts, other input data types (e.g., RNA, Germline_Genotype, Fusion, etc.) could potentially offer an increased range of drug options for patients (**Figs. 4b-d**). Moreover, the drug response module of POI was validated in the FUSCC cohort, where it suggested an additional 55 types of drugs in addition to prioritizations from other modules (**Fig. 4e**). This further confirms POI’s capability to propose a more comprehensive set of drugs, including targeted therapy and chemotherapy. Collectively, these results emphasize the promise of POI in delivering comprehensive precision drug prioritizations by parsing multiomics data, offering the possibility of treatment for a wider range of patients.

### Enhancing anti-cancer drug prioritization through indirect evidence prediction

When a patient’s genomic and transcriptomic variants cannot directly match clinical evidence conclusions in PreMedKB, POI employs a strategy of leveraging biological associations within pathways to identify indirect evidence and provide prioritization for patients who do not match a targeted drug.

Specifically, POI begins by annotating the patient’s germline and somatic SNVs/Indels, taking into account indicators such as population frequency, predicted deleteriousness of variants, and clinical significance to identify potentially abnormal genes. POI then examines their enrichment in the same pathway as the gene corresponding to the actionable therapeutic alteration, utilizing Hallmark gene pathway information [27]. If the aberrant genes belong to the same pathway and show a high correlation (protein-protein interaction score acquired from the STRING database [28] > 0.99), the corresponding drugs are considered potentially effective, and the evidence level for these inferred drugs is assigned as level E.

To assess the effectiveness of indirect evidences provided by POI, we conducted a statistical analysis of drug prioritizations in the cohorts and observed that inferred drugs derived from POI exhibit promising potential in preclinical studies. Taking TCGA’s ovarian cancer patients as an illustrative example, within the test cohort, approximately 71% of patients were eligible for direct drug prioritizations, while 21% were eligible for inferred drug prioritizations, derived from the POI indirect evidence module. The inference process primarily relies on the TP53 mutation status identified by POI. It is important to note that the existing database does not provide specific prioritized drugs for TP53 mutations in ovarian cancer patients. Nevertheless, leveraging pathway associations, POI established a connection between the TP53 and BRCA1 genes [29]. Remarkably, for ovarian cancer patients harboring BRCA1 germline or somatic mutations, both the FDA and NCCN guidelines offered corresponding drug prioritizations, such as Olaparib. Consequently, approximately 21% (124 individuals) of TCGA ovarian cancer patients can receive the tailored treatment advice (**Fig. S4**). Moreover, preclinical investigations have already demonstrated the inhibitory effects of POI-inferred drugs on the growth of xenograft tumors derived from ovarian cancer patients with wild-type ATM and TP53 mutant backgrounds [30]. These findings provided additional support for the potential of POI-inferred drugs, underscoring its promise in clinical applications.

### Web-based interface

POI is a user-friendly web server whose workflow consists of three main pages, including a *Query* Page (input), an *Intermediate* Page (submission and analysis), and a *Report* Page (output) where the therapeutic details can be viewed by clicking the row of the tables in the therapeutic overview (**Fig. 5**).

**Fig. 5.**
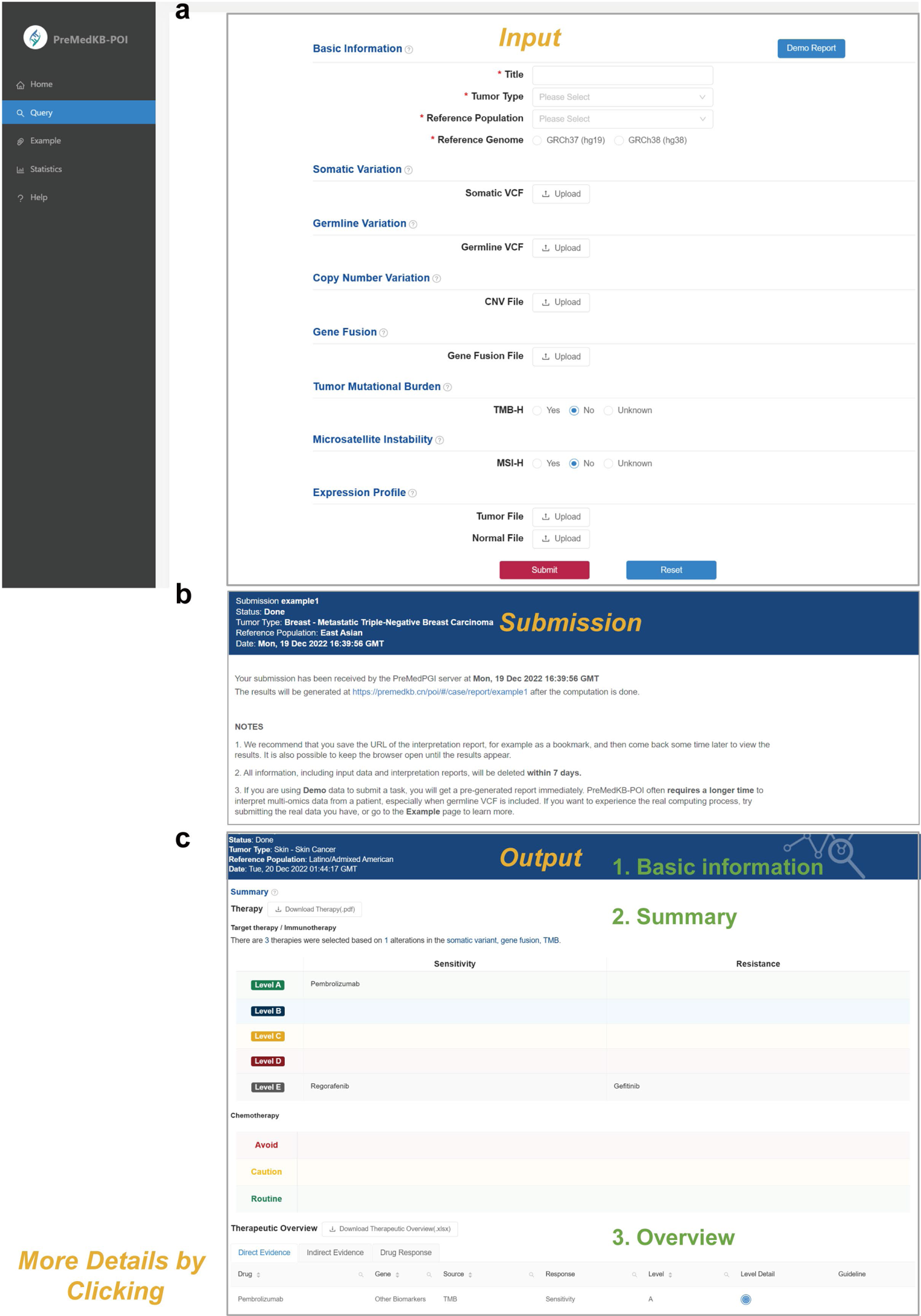
Interface of POI web server. The workflow and output of POI web server. **a** The *Query* Page allows users to input clinical information and personal omics data in different modules. The *Intermediate* Page shows the report address and notes after submission. **b** The *Report* Page displays the basic information of the case, a summary of drug recommendations in two tables, and a therapeutic overview of each drug in three tabs.

In the *Query* Page, the user can select clinical information and upload his/her omics data in each module according to the “*?”* tips (**Fig. 5a**). Upon clicking the *Submit* button, the user is directed to the *Intermediate* Page, where the report address and associated notes are prompted. (**Fig. 5b**). Once the computation is complete, the *Report* Page can be accessed via the provided link. At the top of the *Report* Page, the basic case information can be viewed, followed by two tables summarizing the drug prioritizations. The therapeutic overview of each drug is presented in three tabs: “*Direct Evidence*”, “*Indirect Evidence*”, and “*Drug Response*”. (**Fig. 5c**). Further details of a specific drug’s therapeutic and biomarker information can be explored by selecting the corresponding row in the therapeutic overview. Additionally, the content of the biomarker detail section includes a gene expression distribution for comprehensive analysis.

To assist users in understanding the report generation process, POI offers a Demo Report feature, pre-filled with relevant data. Furthermore, three POI report examples with test data are available, providing users with a better grasp of the POI report structure.

### Use case

We provide an example from a Chinese woman patient from the FUSCC cohort who had metastatic triple-negative breast carcinoma to showcase the effectiveness of the POI in identifying and interpreting actionable therapeutic alterations from individual patients’ multiomics profiles. The detailed reports can be accessed in **Example 1** on the website (https://premedkb.cn/poi/#/case/report/example1).

Specifically, the patient’s somatic Variant Call Format (VCF) data, germline VCF, CNV, and gene expression files, which were fed to POI, were obtained from our previous study [23, 24]. The POI result report reveals the identification of 26 actionable therapeutic alterations in this particular case, accompanied by corresponding 54 drug prioritizations (therapies) varying from level A to level E.

Within the “*Direct Evidence*” tab of the report, the combination therapy of Alpelisib and Fulvestrant with evidence Level A was recommended for this patient with PIK3CA mutation, Lapatinib, and Patritumab Deruxtecan with evidence Level B based on the overexpression of ERBB3 gene from gene expression file was recommended. Additionally, there were several drugs, with evidence of Levels C or D, based on distinct mutations (somatic mutation and CNV) in other genes, as indicated by the somatic VCF data.

In the “*Indirect Evidence*” tab, POI employed inference strategies to prioritize drugs based on the association between the genes BRCA1, CDKN2A, and RB1 with key cancer gene TP53 in the E2F Targets and P53 Pathway through inference strategies of POI were displayed as assigned evidence Level E.

Furthermore, in the “*Drug Response*” tab, the drug response and related phenotypes were predicted based on the resolved diplotype of the patient from her germline VCF. The chemotherapy drugs were summarized in the table of chemotherapy by dividing them into three categories, including avoid use, use with caution, and routine use.

## Discussion

In the domain of precision oncology, the quest for personalized therapy is predicated on the unique molecular signatures of individual patients [31]. Yet, the endeavor to pinpoint optimal treatments is hampered by disparities across oncological knowledgebases, the constraints of manual interpretation, and the insufficient harnessing of genomic data [3]. Despite previous efforts recognizing the importance of including non-somatic variations and adopting multifaceted analytical approaches [12–15], there is still a lack of a comprehensive tool that can prioritize anti-cancer drugs by integrating genomic and transcriptomic data at the individual level. Therefore, we developed POI based on the foundation of PreMedKB [32], a user-friendly system that integrates disease, gene, variant, drug, and clinical evidence information from multiple databases. POI utilizes a harmonized knowledge network and a multi-dimensional interpretation strategy to provide comprehensive drug prioritization. It prioritizes targeted and immunological drugs based on somatic variants, genomics signatures, pathogenic germline variants, and aberrantly expressed genes, thereby aiding in precise treatment selection. Additionally, POI provides repurposing drugs for patients without actionable therapeutic alterations by considering the association of aberrant alterations in specific biological pathways. By integrating comprehensive knowledge, resolving germline diplotypes, and providing chemotherapy drugs with phenotype prediction, POI enhances the clinical utility of precision oncology. The validation of POI on diverse datasets confirms its effectiveness in identifying actionable therapeutic alterations and expands access to therapeutics.

We compared POI (https://premedkb.cn/poi/#/) with five other tools for precision drug prioritization based on omics data of individual patients: PORI [12], MOAlmanac [13], CCAS [33], PanDrugs [15, 34], and Cancer Genome Interpreter (CGI) [8]. We demonstrated that POI outperforms other tools in several aspects, such as input data types, output formats, knowledgebases, evidence levels, drug response prediction, pathway inference, and off-label use cases. POI has three main advantages over other tools. First, POI can analyze a wide range of data types that cover both somatic and germline variants, as well as expression profiles and clinical information. POI can perform tumor-normal paired analysis for more accurate variant calling and expression profiling. This allows POI to identify more comprehensive actionable therapeutic alterations than other tools. Second, POI integrates multiple databases, standardizes knowledge from different sources, and can prioritize potential drugs that do not have direct evidence in existing databases through biological pathway inference. Third, POI provides a comprehensive report that includes drug prioritizations based on different evidence levels from multiple sources, which could help clinicians decide whether patients should use chemotherapy drugs or targeted drugs. We have summarized the detailed comparison results in **Table 1**.

**Table 1.** Comparison of online interpretation tools for drug prioritization.

| Type | Resource | <u>POI</u> | PORI [12] | MOAlmanac [13] | CCAS [14] | PanDrugs [7, 15] | CGI [8] |
| --- | --- | --- | --- | --- | --- | --- | --- |
| **Variant** | SNV/ INDEL | ✓ | ✓ | ✓ | ✓ | ✓ | ✓ |
|  | CNV | ✓ | ✓ | ✓ | ✓ | ✓ | ✓ |
|  | Fusion | ✓ | ✓ | ✓ |  |  | ✓ |
|  | RNA expression | ✓ | ✓ |  | ✓ | ✓ |  |
|  | Germline variant<sup>a</sup> | ✓ |  | ✓ |  | ✓ |  |
| **Application** | Clinical evidence | ✓ | ✓ | ✓ |  |  | ✓ |
|  | Drug response | ✓ |  |  |  | ✓ | ✓ |
|  | Drug repurposing | ✓ | ✓ |  |  | ✓ | ✓ |
|  | Interaction visualization<sup>b</sup> | ✓ | ✓ | ✓ | ✓ | ✓ |  |
| **Knowledgebase** | Term normalization | ✓ | ✓ |  | ✓ | ✓ |  |
|  | Database integration<sup>c</sup> | • CGI<br>• CIViC<br>• COSMIC<br>• My Cancer Genome<br>• OncoKB<br>• PharmGKB | • CGI<br>• CIViC<br>• COSMIC<br>• DoCM<br>• OncoKB | • TARGET<br>• COSMIC | • COSMIC<br>• DiseaseMeth<br>• Disease OncoKB<br>• DoCM<br>• CGP | • CGI<br>• CIViC<br>• COSMIC<br>• My Cancer Genome<br>• OncoKB<br>• PharmGKB<br>• TARGET | • CIViC<br>• DoCM<br>• OncoKB |

<sup>a</sup> Contains both direct recommendations based on mutations, and filtering based on germline genotypes to determine drug responses.
<sup>b</sup> e.g., pathway, data of cancer cohort.
<sup>c</sup> Databases that fit this category should contain the relationship between variants and therapeutic evidence.

Besides, the integration of different knowledge bases significantly enhances the overall knowledge coverage, leading to the identification of a more extensive range of actionable therapeutic alterations and facilitating the prioritization of a greater number of drugs for individual patients. The validation of database integration using three cancer datasets demonstrated that the annotation results for drug prioritization in patients not only relied on commonly shared drugs across all integrated knowledge bases but also encompassed specific drugs unique to individual or two specific knowledge bases. This observation further highlights the comprehensive nature of treatment-related knowledge within the PreMedKB database (**Fig. S5**). However, it is important to note that variations in terminology formulation and the inclusion of rare variants within a single database contribute to the distinctive features observed among databases (**Fig. 2c**). This variation may be magnified as the reliability of clinical evidence decreases, emphasizing the need for careful consideration and evaluation of the available knowledge sources.

Although POI can expand treatment options for patients through more comprehensive actionable therapeutic alterations, its effectiveness needs to be further verified. Direct application of drug prioritizations by POI and other similar online portals to clinical practice is not always possible. Clinical interventions require review by, for example, the Molecular Tumor Board (MTB) and similar bodies, and better integration of the system with the practice of clinical oncologists is necessary [35]. Additionally, the adoption of variant information exchange standards by the community is not complete, and the Association for Molecular Pathology, American Society of Clinical Oncology, and College of American Pathologists (AMP/ASCO/CAP) classification standards used in this study have room for refinement and are only applied by about 70% of investigators [36]. Furthermore, systematic interpretation of results requires rigorous validation of analytical and clinical validity in combination with assay reagents, with the ultimate focus being on clinical utility for precision medicine [37, 38].

Looking ahead, the ongoing development and refinement of POI could utilize an open-source community development model [39], promising to advance personalized therapy and improve patient outcomes. By integrating additional omics data, such as proteomics [40] and methylation [14] data, POI can continue to evolve as a valuable tool that expands access to therapeutics and enhances the precision of treatment decisions.

## Conclusions

POI serves as a valuable tool for the comprehensive prioritization of anti-cancer drugs by effectively identifying actionable therapeutic alterations in a patient’s multiomics profiles, thereby broadening the availability of targeted therapeutics. The webserver of POI assists researchers and clinicians in navigating the complexities of these profiles, facilitating precise therapy decision-making. Notably, the capability of POI to prioritize drugs based on robust preclinical evidence highlights its potential to provide substantial benefits to patients, particularly in off-label use cases. Ongoing efforts focused on the continuous development and refinement of POI hold great promise for advancing precision medicine and ultimately enhancing patient outcomes.

## Supporting information

Additional file 1 POI Supplementary figures

## Methods

### Construction of knowledgebase

We used our previously reported knowledgebase PreMedKB [32] to provide clinical and biological evidences for interpretation. For facilitating cancer genomics interpretation, most databases have been updated and cancer-related databases were added in PreMedKB. Here, we briefly described the construction and updating of PreMedKB.

### Data sources

PreMedKB serves as the foundation for the POI algorithm and consists of two distinct layers: the meta-knowledgebase layer and the domain knowledgebase layer. The meta-knowledgebase layer encompasses databases related to diseases, genes, variants, and drugs, along with their metadata, including names, synonyms, functions, *etc*. The domain knowledgebases established relationships among these elements and served as data sources that provided insights into the clinical significance of diseases, genes, variants, and drugs. Notably, the entries within these knowledgebases represented connections between two or more of the elements mentioned above.

To ensure a comprehensive integration of clinical evidence conclusions on cancer genomics, the updated version of PreMedKB incorporated reliable data obtained from expertly curated databases, as summarized in **Table S1**. These data sources were managed as independent databases using MySQL. The construction process involved the design of the entity-relationship diagram for the database, formulation of the data dictionary, implementation of data preprocessing techniques, and subsequent data importation. These coordination strategies closely resembled those employed in the original version of PreMedKB, with the primary distinction lying in the assimilation of updated data sources.

### Meta database construction

To enable interoperability and establish connections between research and clinical settings, PreMedKB provides a rich vocabulary in its metadata databases. Standard names and synonyms are retrieved from various data sources. Gene names are standardized using the HUGO Gene Nomenclature Committee (HGNC, https://www.genenames.org/) [41], NCBI Entrez Gene (https://www.ncbi.nlm.nih.gov/gene/) [42], and Ensembl. Variation information was obtained primarily from dbSNP (https://www.ncbi.nlm.nih.gov/snp/) [43], Clinical Variation Database (ClinVar, https://www.ncbi.nlm.nih.gov/clinvar/) [44], Catalogue of Somatic Mutations in Cancer (COSMIC, https://cancer.sanger.ac.uk/cosmic) [45], and gnomAD’s calculated allele frequencies for population frequency annotation. Additionally, ANNOVAR [46] was employed for annotating the functional effects of variations. Disease metadata, including name, definition, and ontology structure, were harmonized using OncoTree (http://oncotree.mskcc.org/) [47], Mesh (https://meshb-prev.nlm.nih.gov/search) [48], and the NCI Thesaurus (NCIt, https://ncithesaurus.nci.nih.gov/ncitbrowser/) [49, 50]. Drug-related information, encompassing structure, pharmacology, pharmacogenomics, clinical stages, and product details, were integrated using resources such as NCIt, ChEBI (https://www.ebi.ac.uk/chebi/) [51], ChEMBL (https://www.ebi.ac.uk/chembl/) [52], and DrugBank (https://go.drugbank.com/) [53].

### Domain knowledgebase integration

Domain knowledgebase integration primarily involves three dimensions of information: clinical evidence conclusions, the landscape of the genome and transcriptome profiles, and biological pathway knowledge. Prominent knowledgebases such as OncoKB [54], CIViC [39], My Cancer Genome (MCG) [55], CGI [8], PharmGKB [56, 57], and NCCN Drug (https://www.nccn.org/#) provided clinical evidence conclusions regarding target therapies, immunotherapies, chemotherapies, pharmacogenomics, and pathogenic sites in cancer. The landscape of the genome and transcriptome profiles was derived from The Cancer Genome Atlas (TCGA, https://cancergenome.nih.gov/), a valuable resource encompassing diverse integrative cancer genomics data from various human tissues. In this study, we utilized the RNA expression landscape of TCGA [58] to construct a RNA reference database of gene expression levels, facilitating the assessment of abnormal gene expression in patients. Additionally, MsigDB [27] offered biological pathway knowledge associated with hallmark genes, providing an additional dimension for linking existing clinical evidence with patients’ omics profiles.

The meta-knowledgebases were constructed with comprehensive lexicons for the four main elements: diseases, genes, variants, and drugs. This enabled the matching of nodes in the domain knowledgebases with meta-IDs using lexical matching. To ensure accuracy and consistency, duplicate semantic relationships were eliminated, and nodes were assigned higher confidence ratings accordingly.

### Normalization of clinical evidence levels

Given the inherent variability in describing evidence levels within different knowledgebases, a harmonization process involving manual mapping is necessary to establish a unified standard.

It was worth noting that the release of the AMP/ASCO/CAP somatic classification guidelines [59] took place subsequent to the design of the Variant Interpretation for Cancer Consortium (VICC) knowledgebases and was partially influenced by them. While the evidence levels within the knowledgebases exhibit compatibility with the AMP/ASCO/CAP guidelines [59], it was important to acknowledge that they were not entirely identical. Consequently, a comprehensive mapping of the evidence levels provided by each knowledgebase was conducted to align them with the AMP/ASCO/CAP guidelines. For instance, Level A refers to biomarkers that predict response or resistance to FDA-approved therapies or professional guidelines for a specific type of tumor [59]. Detailed descriptions and mappings of each evidence level can be found in **Table 2**.

**Table 2.**
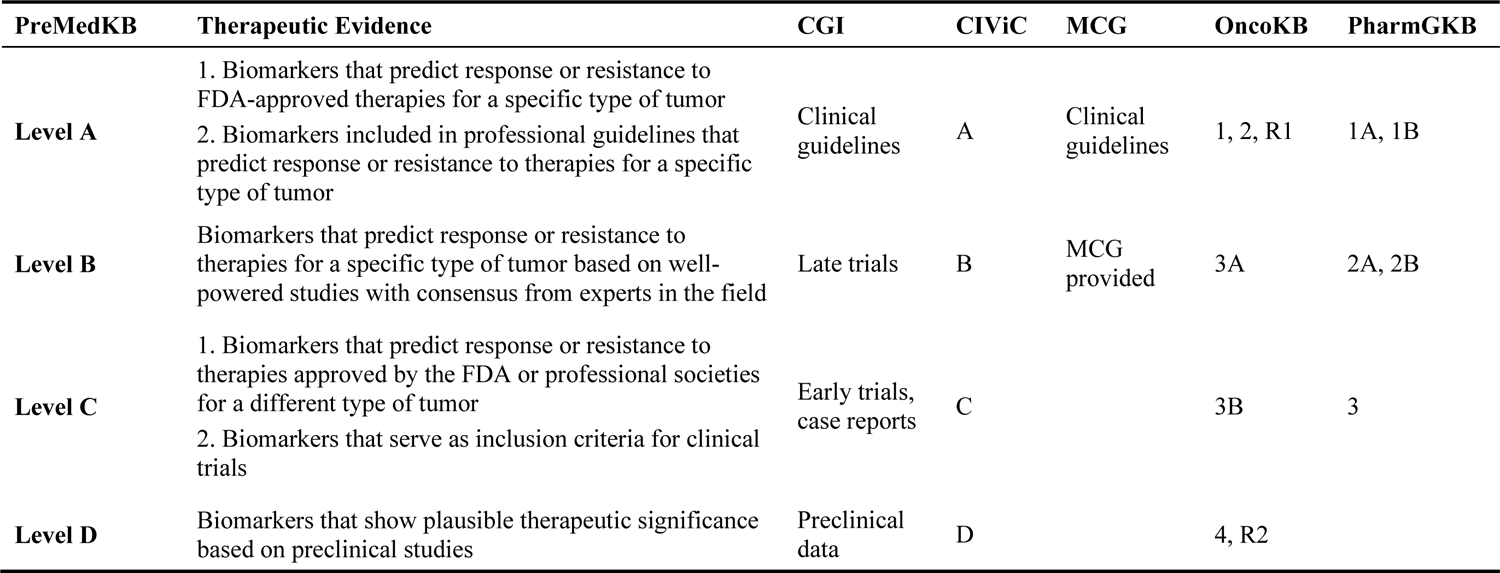
Harmonizing knowledgebase-specific evidence levels based on AMP/ASCO/CAP guidelines. [59].

### Analysis modules and drug prioritization strategies

The analysis module of the POI system consists of three components, based on direct evidence, indirect evidence, and drug response. Genomic and transcriptomic data submitted by users undergo preprocessing and normalization by the POI system. Subsequently, the ANNOVAR tool is employed for variant annotation in VCF files, thereby enhancing the subsequent interpretation process. The clinical information provided by the users, including tumor type, plays a crucial role in facilitating the accurate prioritization of targeted and immunotherapeutic drugs, while reference population information will be utilized to predict the response to chemotherapy drugs and assess the associated phenotypes. These three modules collectively identify and interpret the actionable therapeutic alterations, providing comprehensive and precise guidance for the prioritization of targeted, immunological, and chemotherapeutic drugs.

### Direct evidence

Direct evidence pertains to the utilization of clinical and experimental research findings specific to genetic variants associated with particular tumor types and treatments. This evidence is sourced from the integrated domain knowledgebase of PreMedKB (**Table S1**). POI can identify the actionable therapeutic alterations through the analysis of various patient files, including somatic and germline SNV/Indel, CNV, gene fusions, TMB, MSI, and gene expression data, enabling effective prioritization of targeted and immunological drugs.

When handling annotated somatic and germline SNV/Indel, POI employs the following strategy to match them with clinical evidence in PreMedKB. Direct matching is employed for variants with well-defined amino acids or bases. If the clinical evidence specifies variants only for a specific mutation class in a gene or on a specific exon, direct matching is performed against those known conditions. For instance, in the case of variants in the truncated mutation category, matching is based on the annotated variant category (*e.g.*, stopgain, frameshift insertion, frameshift deletion, frameshift block substitution). In situations where clinical evidence refers to oncogenic mutations without detailed variant information, POI first determines if the patient has a variant considered oncogenic in PreMedKB (integrated across databases) and then determines if it is annotated as pathogenic or likely pathogenic by ClinVar. Wild-type variants, such as *KRAS* and *NRAS* genes, are assessed separately after resolving other variants.

Genomic signatures, such as TMB and MSI, play a significant role in individual tumor genomes and have implications in oncology treatment. However, given the lack of standardized calculation methods for TMB and MSI, POI offers options of high tumor mutation burden (TMB-H) and high microsatellite instability (MSI-H) on the website rather than performing direct calculations based on user-submitted VCF files. For CNV, POI resolves the gene status in the CNV file, including “gain”, “loss”, or “neutral”. Regarding gene fusions, POI parses the gene pairs provided in the file.

To effectively utilize the patient’s transcriptomic data, POI converts raw read counts into Counts Per Million (CPM). Genes with at least 10 counts and a minimum CPM of 0.5 in both tumor and normal tissues are considered expressed genes, which are used for further analysis. A log2 transformation is applied to the CPM values, with an additional value of 0.01 added to the CPM of each gene to avoid infinite values. A gene is considered as under-expressed if the relative expression of in tumor versus normal tissues (fold change of tumor/normal, log2 transformed) is ≤ −3.5, while a gene is considered as over-expressed if its relative expression is ≥ 3.5. The threshold was established and validated using a breast cancer cohort, with the expression status of *HER2*/*ERBB2* serving as the ground truth (**Fig. S6** and **Table S2**). Additionally, POI calculates the patient’s gene expression in proportion to the TCGA cohort of the same cancer type to further confirm the abnormal expressed gene for drug prioritization.

In cases where disease groups share the same ancestry or descent, they are assigned the same clinical evidence, and drug prioritization is directly assigned according to the evidence level normalized by POI. However, if a patient possesses a variant recognized by professional guidelines (Level A), but their cancer type does not share ancestry or descent with the supported indication in the clinical evidence, the drug prioritization is downgraded from Level A to Level C.

### Indirect evidence

POI begins by annotating the patient’s germline and somatic SNVs/Indels, taking into account indicators such as population frequency, predicted deleteriousness of variants, and clinical significance to identify potentially abnormal genes. Variants that meet any two of the three following indicators are identified as potentially aberrant variants or genes. Firstly, rare variants with a frequency of less than 1/1000 in the population, indicating low prevalence in the general population and potential association with specific diseases or genetic disorders [60, 61]. Secondly, variants annotated as pathogenic or likely pathogenic in ClinVar [44] are considered. Lastly, variants are classified as pathogenic mutations affecting protein function if at least one of the indicators (SIFT, LRT, MutationTaster, MutationAssessor, FATHMM, PROVEAN, MetaSVM, MetaLR) designates them as such.

POI then examines their enrichment in the same pathway as the gene corresponding to the actionable therapeutic alteration, utilizing Hallmark gene pathway information [27]. If the aberrant genes belong to the same pathway and show a high correlation (protein-protein interaction score acquired from STRING database [28] > 0.99), the corresponding drugs are considered potentially effective, and the evidence level for these inferred drugs is assigned as level E.

### Drug response

Chemotherapeutic drugs are known for their broad-spectrum efficacy against diverse tumor cell types, making them a crucial component of cancer treatment [62]. Understanding the influence of genetic variations on drug response holds significant potential for refining chemotherapy protocols, mitigating adverse effects, and enhancing therapeutic outcomes for individuals with cancer [63]. Hence, the analysis of germline variation data, particularly gene polymorphisms associated with drug metabolism and drug targets, enables the prediction of patient sensitivity to chemotherapeutic drugs to improve the precision medicine. To accomplish this, our module incorporates our previously developed pharmacogenomics annotation tool *PAnno* [25] into the POI framework. Through the analysis of germline variants, we infer the genotypes/diplotypes of genes related to chemotherapeutic drugs, enabling the prediction of patient phenotypes about toxicity, dosage, efficacy, and drug metabolism. Finally, drugs are categorized into three levels: avoid caution, and routine.

### Multiomics data processing of three external cohorts

#### TCGA

The TCGA dataset, consisting of genomics and transcriptomics data from 11,005 patients, was downloaded from cBioPortal [21] (https://www.cbioportal.org/), which includes all studies from *_tcga_pan_can_atlas_2018: ACC, BLCA, BRCA, CESC, CHOL, COADREAD, DLBC, ESCA, GBM, HNSC, KICH, KIRC, KIRP, LAML, LGG, LIHC, LUAD, LUSC, MESO, OV, PAAD, PCPG, PRAD, SARC, SKCM, STAD, TGCT, THCA, THYM, UCEC, UCS, and UVM. The TCGA cohort covered 33 cancer types, normalized by 29 MSKCC tissue codes (**Table S4**). The data_mutaions files were converted to VCF files using the maf2vcf tools (https://github.com/mskcc/vcf2maf) developed by MSKCC, and CNA data files were binned into individual CNV files by patient ID, with the gene status divided by the same thresholds used in MSK-IMPACT above. TCGA fusion gene datasets were obtained from ChimerDB 4.0 (http://www.kobic.re.kr/chimerdb/) [64], and the MSI datasets were downloaded from the TCGA MSI landscape [65] with MANTIS scores [66] for identifying MSI-H patients (where MSI-H was defined as MANTIS score > 0.4). To identify “TMB-H” patients, the TMB value was calculated based on the mutant allele frequency (MAF) files downloaded through the R package “TCGAmutations” [67]. Furthermore, clinical information and gene expression files were collected through the R package “TCGAbiolinks” [68] for subsequent analysis.

#### MSK-IMPACT

The MSK-IMPACT dataset is a genomics repository of 10,945 patients, which was obtained from cBioPortal (http://cbioportal.org/msk-impact) [21] and a previous study [26]. The raw data was subsequently partitioned into five categories: somatic mutation, CNV, fusion gene, MSI-H, and TMB-H data (**Table S3**). The conversion of the data_mutaions file to VCF files for each individual patient was performed utilizing the maf2vcf tools developed by MSKCC, available on GitHub (https://github.com/mskcc/vcf2maf). The CNV files were annotated such that genes with CNA count greater than 2 were labeled as “gain”, those less than −2 were labeled as “loss”, otherwise were labeled as “neutral” [69]. The MSISensor scores [70] of the MSK-IMPACT dataset were used to identify “MSI-H” patients (where MSI-H was defined as MSISensor score > 10) [71], while the TMB was used to identify “TMB-H” patients (where TMB-H was defined as ≥10 mutations/Mb according to FoundationOne CDx (F1CDx) [72]). Finally, all data files were grouped by unique patient ID for subsequent analysis.

#### FUSCC

In this study, we utilized the FUSCC dataset [23], which consists of 427 patients and integrates various types of omics data, including germline and somatic variations, CNVs, and tumor-normal paired RNA expression profiles (**Table S5**). Specifically, the germline variations of 279 patients were obtained through in-house pipelines and were previously unpublished. The purpose of including these germline variations was to enable comprehensive performance validation of the multiomics data analysis. To identify patients with TMB-H, we calculated TMB values based on the MAF files that were obtained from figshare (http://dx.doi.org/10.6084/m9.figshare.19783498.v5).

#### Statistical analysis and validation of external cohorts

We performed a comprehensive statistical analysis of the validation results from three external cohorts. The proportion of patients assigned drug prioritization according to the highest level of evidence was independently calculated for each cohort. The results were visualized using stacked bar charts created with the R package ggpubr v0.6.0. Furthermore, a detailed statistical analysis was conducted to examine the different alterations observed in the drug sources within each cohort. The findings of this analysis were effectively presented using an upset plot, utilizing the R package ComplexUpset v1.3.3. Additionally, a specific investigation was carried out on the core modules of the POI system for drug sources within the FUSCC cohort. The results of this investigation were visualized through a Venn diagram created using the R package eulerr v7.0.0.

#### Webserver construction

POI was employed various technologies in its front-end user interface, including the React framework (https://reactjs.org/), Ant Design (https://ant.design/), and Apache Echarts (https://echarts.apache.org/). The last technology was primarily employed in the *Statistics Page* to enable the visualization of large amounts of data. In the back-end architecture, the Flask-based (https://flask.palletsprojects.com/) web framework was used to receive and process user requests, while also facilitating communication between the front-end interface and the underlying database. The REST architecture style was utilized in the development of POI to reduce the intricacy of development and enhance system scalability. MySQL database management system was utilized to store and manage all data within the system.

#### Availability and requirements

Project name: POI

Project home page: https://premedkb.cn/poi/#/homepage

Operating system: Platform-independent

Programming language: Python, MySQL

Other requirements: R version greater than 3.5

License: Crick Non-commercial License Agreement v2.0

Any restrictions on use by non-academics: Commercial use will require a license from the rights holder. For further information, contact.

## Declarations

### Ethics approval and consent to participate

Not applicable.

### Consent for publication

Not applicable.

### Availability of data and materials

The tool is freely available through the API or the web interface at https://premedkb.cn/poi/#/homepage. An updated version of PreMedKB for variant interpretation is included in the software. The germline variant datasets in the FUSCC cohort are not publicly available due to the potential compromise of personal privacy.

## Authors’ contributions

Y.Y., L.G., and Y.L. conceived the study. Y.L. and Q.W.C. developed the POI algorithm. Y.L. and Q.C.C. updated the PreMedKB databases. Q.W.C. collected the data used in the software development and performance validation. Q.W.C. and Y.W. contributed to the validation and interpretation of the results. L.Q.S. developed the webserver. Y.L., Q.W.C., and L.Q.S. drafted the manuscript; L.M.S., Y.Z., L.G., and Y.Y. reviewed it. All authors contributed to the article and approved the final manuscript.

## Acknowledgements

We thank Dr. Xin Hu of Precision Cancer Medicine Center of Fudan University Shanghai Cancer Center for the valuable input in our discussions. We are grateful to CFFF (Computing for the Future at Fudan) and the Human Phenome Data Center of Fudan University for computing support.

## Additional files

### Additional file 1. Supplementary figures

**Figure S1.** Histogram of the number of diseases.

**Figure S2.** Gene expression distribution in the RNA reference database.

**Figure S3.** Overview of the three cohorts.

**Figure S4.** Preclinical studies provide support for inferred drugs from POI.

**Figure S5.** Databases of drug sources for the three external cohorts.

**Figure S6.** Threshold selection for determining the gene status in the RNA module.

### Additional file 2. Supplementary tables

**Table S1.** Data sources of the updated PreMedKB.

**Table S2.** Expression profiles of *ERBB2* gene in FUSCC dataset.

**Table S3.** Cases of the MSK-IMPACT project used in this study.

**Table S4.** Cases of the TCGA project used in this study.

**Table S5.** Cases of the FUSCC project used in this study.

