## Additional file 1 POI Supplementary figures for "A comprehensive personal omics clinical interpreter based on genomic and transcriptomic profiles"

### Additional file 1. Supplementary figures

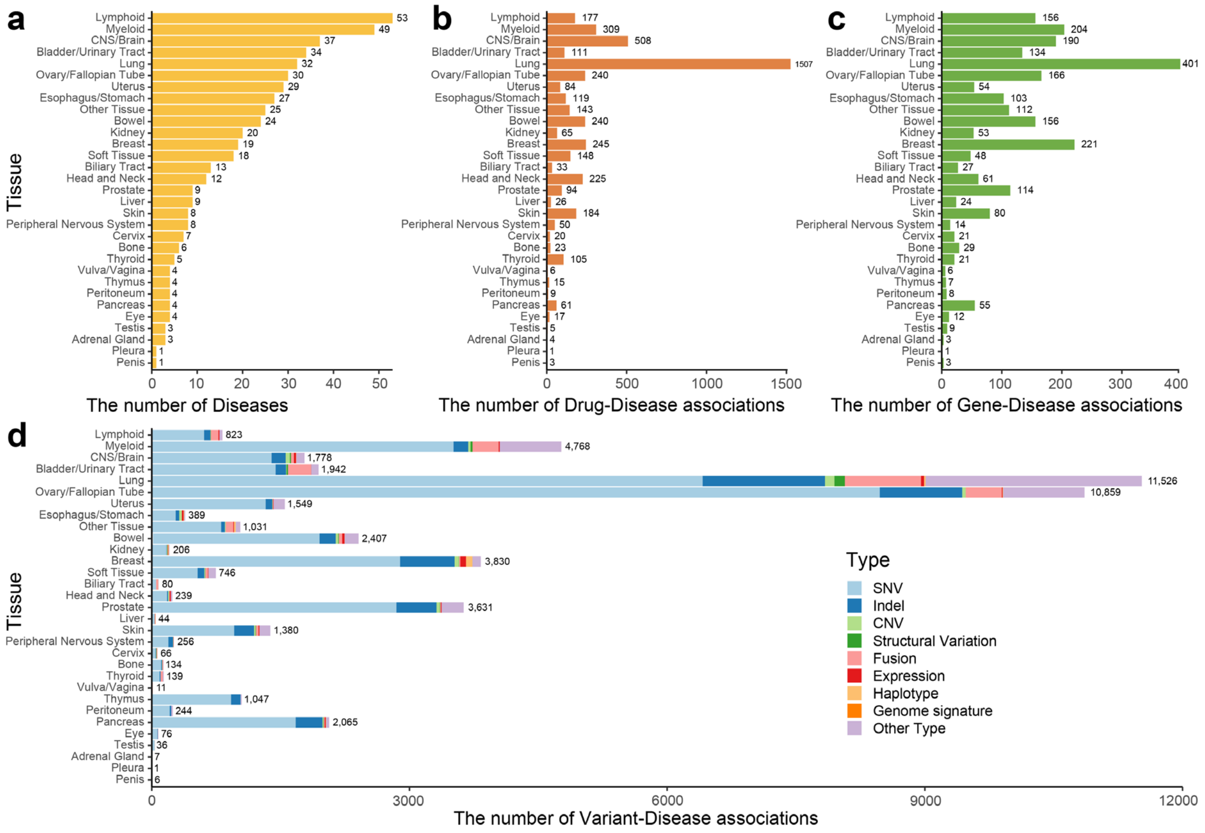


**Figure S1. Histogram of the number of diseases**

The bar plots represent **a** the number of cancer tissue species, **b** drug-disease associations, **c** gene-disease associations, and **d** variants-disease associations across different cancer tissues in the knowledgebase. Each bar represents the frequency of occurrences for the respective association type within the specific cancer tissue.

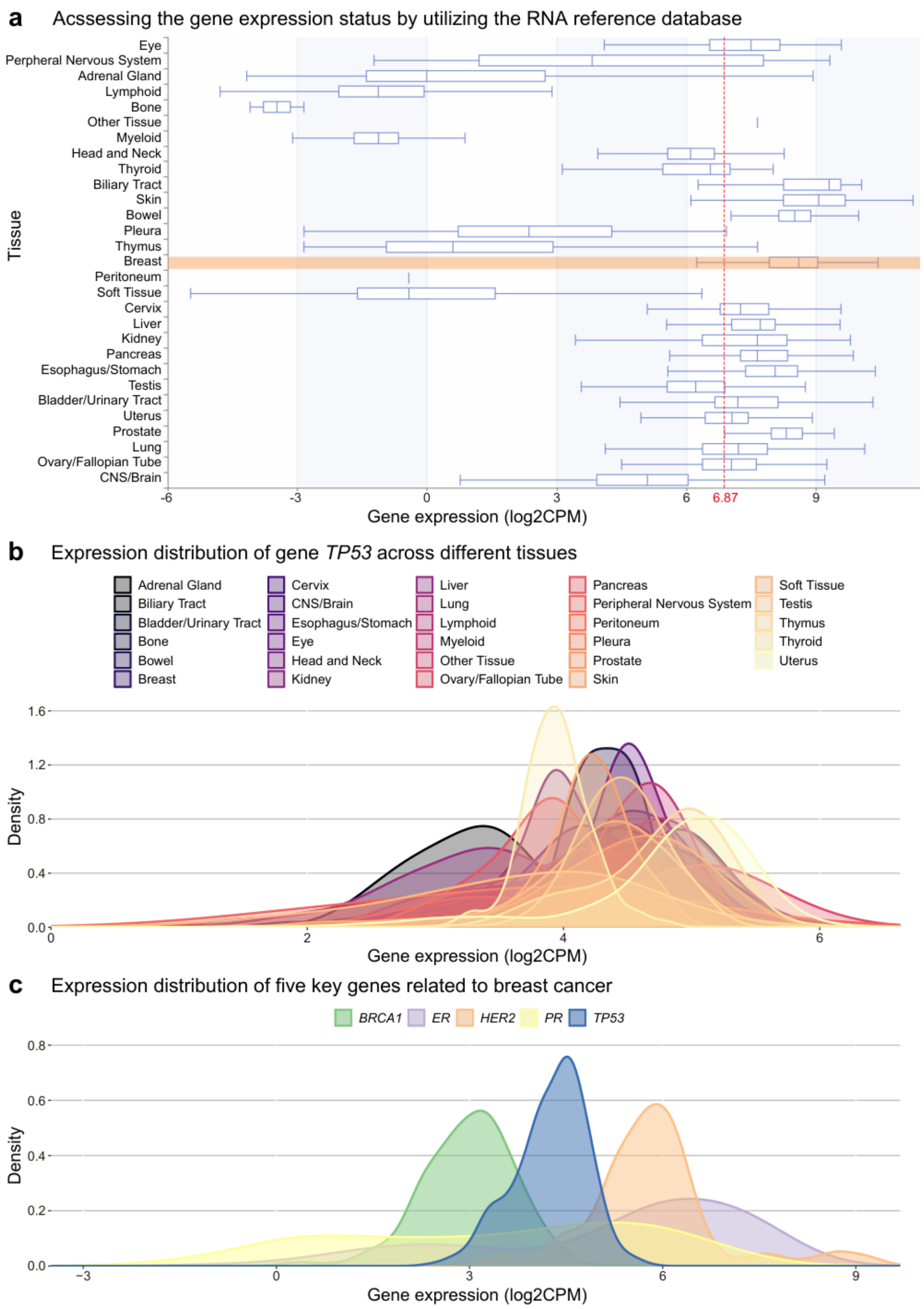


**Figure S2. Gene expression distribution in the RNA reference database**

**a** Boxplot represents an example of utilizing the reference database to assess gene expression of the patient. The gene expression distribution specific to the patient's cancer type in orange, while a red dashed line indicates the position of the patient's gene expression level within the figure. Density plots illustrate **b** the expression distribution of gene *TP53* across different tissues in the RNA reference database and **c** the expression distribution of five key genes associated with breast cancer in the RNA reference database.

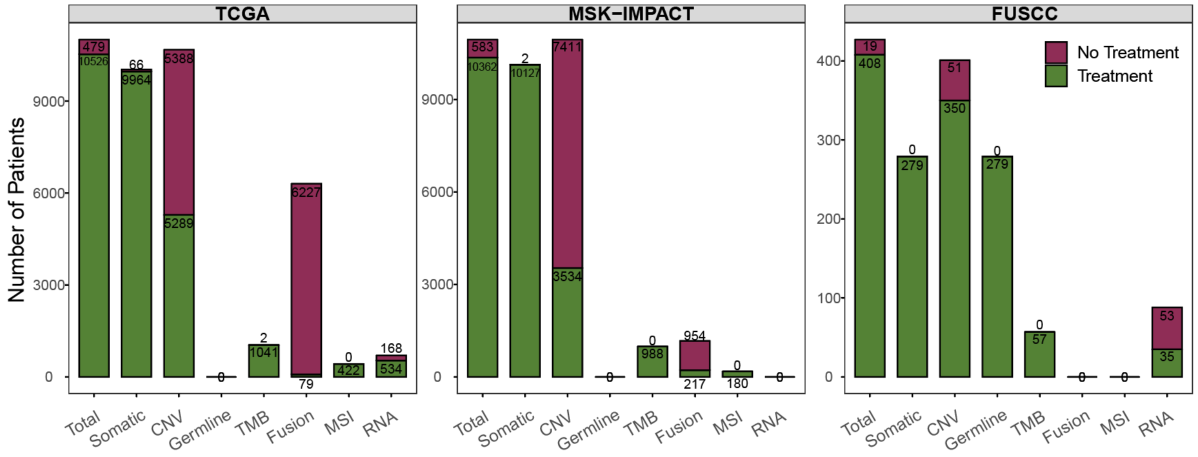


**Figure S3. Overview of the three cohorts**

Bar plots showed the number of patients and the types of multiomics files available for each cohort.

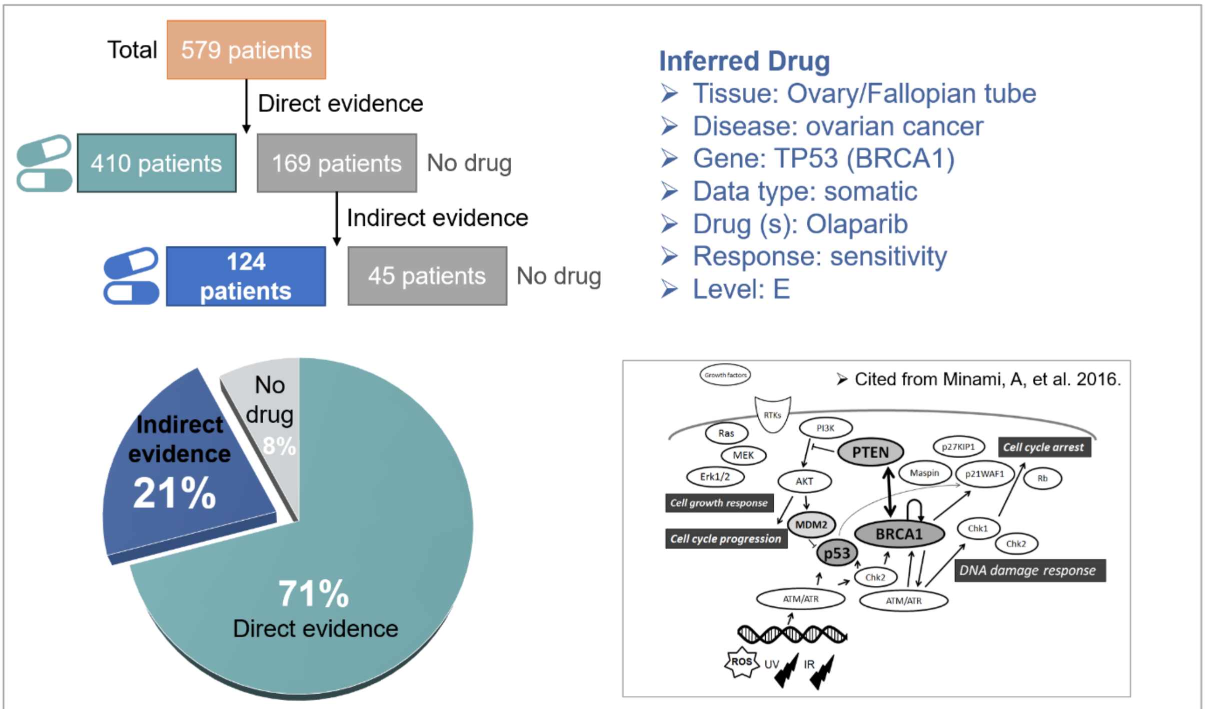


Proportion of drug prioritization module of ovarian cancer patients in the TCGA cohort.

Schematic illustration of the integrative signaling model of tumor suppressors including **p53**, PTEN, and **BRCA1**.

**Figure S4. Preclinical studies provide support for inferred drugs from POI**

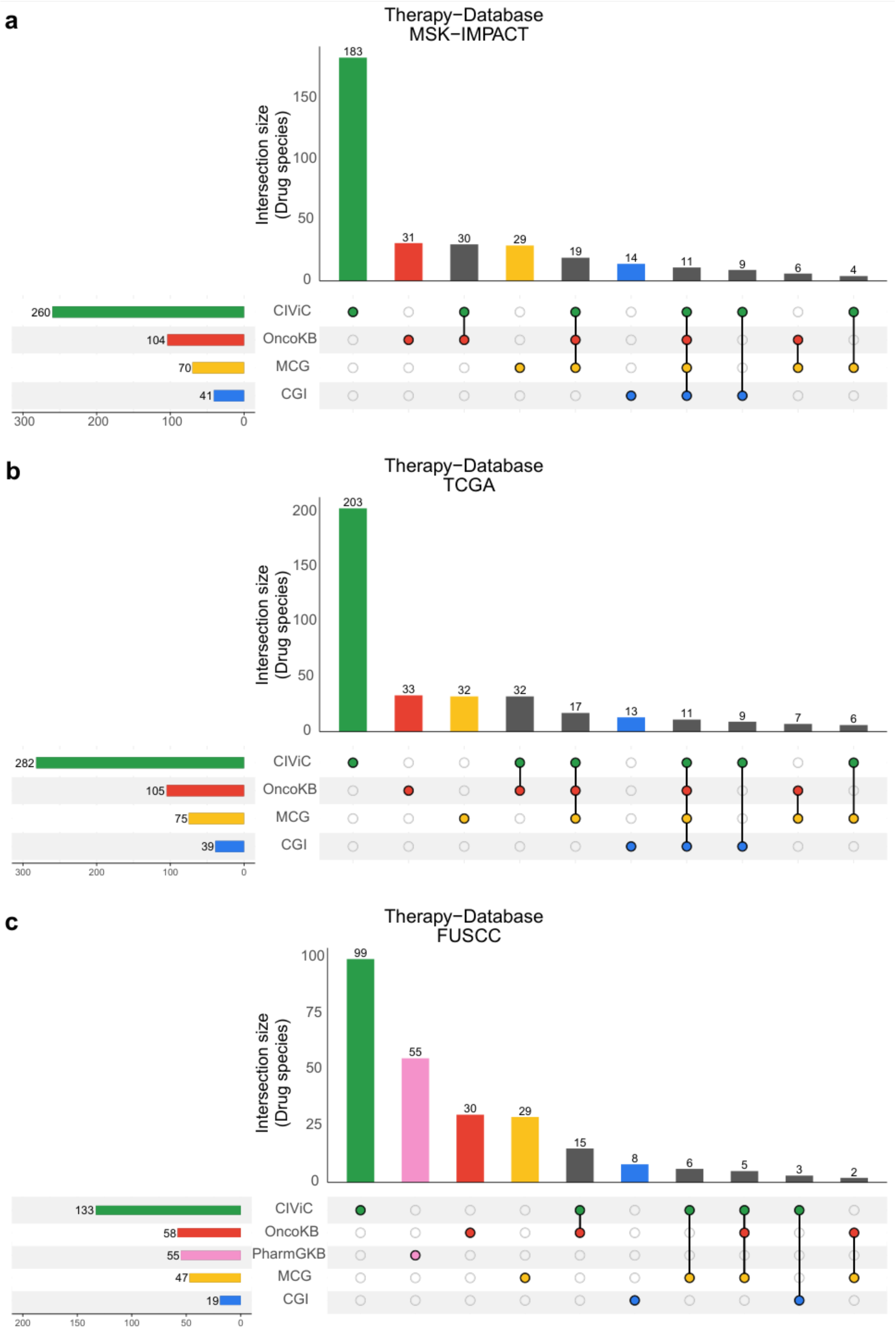


**Figure S5. Databases of drug source for the three external cohorts**

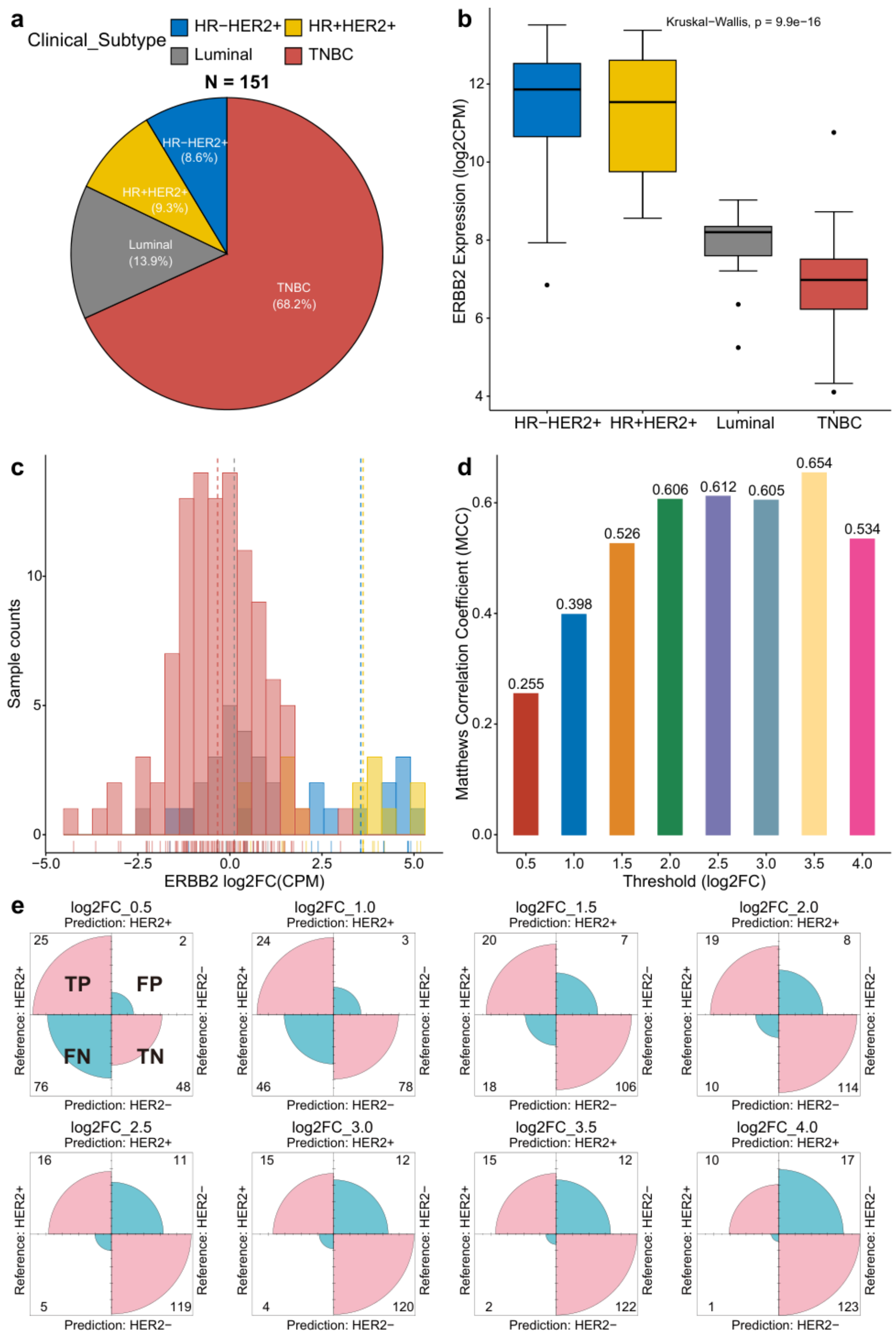


**Figure S6. Threshold selection for determining the gene status in the RNA module.**

**a** Distribution of the breast cancer cohort **b** The expression status of *HER2/ERBB2* gene in different clinical subtypes. **c** The different thresholds (log2 fold change, log2FC) of *ERBB2* gene in different clinical subtypes. **d** The Matthews correlation coefficient

(MCC) corresponding to different thresholds (log2FC) and a higher MCC value indicates more accurate predicted results. **e** The confusion matrix corresponding to different thresholds (log2FC), and the size of the quartet circle area represents the number of samples.
